# HiCPotts: An R/Bioconductor package to identify significant interactions in chromosome conformation capture data and model sources of biases

**DOI:** 10.64898/2026.05.21.726529

**Authors:** Itunu Godwin Osuntoki, Andrew Harrison, Hongsheng Dai, Yanchun Bao, Nicolae Radu Zabet

## Abstract

**Motivation:** Chromosome Conformation Capture methods, including Hi-C, micro-C or Capture-C, are used to map chromatin interactions genome-wide. Most of the existing computational methods do not account for sources of biases (such as DNA accessibility, GC content or TE content) in the data.

**Results:** We previously developed ZipHiC, a Bayesian method based on a the hidden Markov random field (HMRF) model and the Approximate Bayesian Computation (ABC), that uses zero-inflated Poisson distribution to model the noise, signal and false signal of the data and showed that this approach was able to detect biases from DNA accessibility, GC content and TE content in both Hi-C and micro-C data. Here, we present HiCPotts, another Bayesian method based on the HMRF model and the ABC that uses a zero-inflated Negative Binomial distribution instead to model the noise and signal of the data. We systematically show that HiCPotts reduces false positives and increases recovery of true interactions compared to ZipHiC, but also compared to other methods such as FastHiC, Juicer and HiCExplorer. Most importantly, we provide an R/Bioconductor package that allows modelling the noise, signal and false signal using various distributions such as the zero-inflated Negative Binomial (ZINB) and the zero-inflated Poisson distribution (ZIP).

**Availability:** https://bioconductor.org/packages/HiCPotts/

## INTRODUCTION

The genome folds in the nucleus ensuring that distal regulatory elements contact the correct promoters across kilobases to megabase distances (1). Association between distant regions of the genome has been detected by several chromatin capture methods techniques (including Hi-C or micro-C), which allowed building 3D contact maps of the entire genome in individual cells, tissues or developmental stages (2; 3).

Advances in computational methods have progressively improved the spatial resolution of Hi-C and micro-C data (4). Hi-C and micro-C produce large, sparse contact-count matrices and the data can be influenced by technical and biological factors. Fragment length, GC content, and mappability are major Hi-C biases (5). Iterative Correction and Eigenvector decomposition (ICE) balances rows and columns to equalise visibility and remove such biases (6), but assumes all loci are equally observable and becomes computationally expensive at high resolution (6). Knight-Ruiz matrix balancing provides a faster alternative by scaling rows and columns so their sums equal one, yielding a doubly stochastic matrix (7).

More recent methods model biases within explicit statistical frameworks. Hi-C-DC uses distance-stratified negative binomial regression, incorporating GC content, mappability, and fragment length to assess interaction significance (8). Hi-C-DC+, an extension of Hi-C-DC, improves preprocessing, distance-stratified background modelling, and differential analysis, increasing sensitivity and interpretability (9). However, both Hi-C-DC and Hi-C-DC+ treat each interaction independently and ignore spatial dependencies and neighbourhood structure (9).

Other recent approaches include Capricorn, a deep learning framework that super-resolves low-coverage Hi-C matrices to improve loop detection by learning from high-resolution reference maps (10). Capricorn enables signal recovery but assumes that 3D contacts are conserved and could have difficulties identifying novel or cell specific contacts. Furthermore, it does not explicitly model genomic biases or spatial dependencies. AutoHiC uses a deep neural network to jointly learn normalisation weights and correct assembly errors, accounting for biases but still treating contacts independently and omitting spatial correlations (11). HiC2Self, a self-supervised denoising method, improves Hi-C data quality by learning latent structure without labels, but likewise lacks explicit modelling of genomic covariates or neighbourhood dependencies (12). Despite all these advances, most methods still classify contact counts into “signal” versus “background” and treat each contact as independent, neglecting the strong two-dimensional neighbourhood dependencies that characterize contact maps.

One of the ways to address spatial dependence is by using the Bayesian hidden-Markov random-field (HMRF) model. The HMRFBayesHiC method captures spatial dependencies via a hidden Markov random field, offering a principled Bayesian framework that accounts for genomic biases and neighbourhood correlations, therefore making it an early precursor to spatial models like ZipHiC (13; 14). ZipHiC extended on the HMRFBayesHiC by employing the Potts model in modelling spatial dependency directly on the contact matrix (13). It treats each pixel in the Hi-C matrix as part of a 2D lattice and encourages adjacent loci to share similar latent states, enabling borrowing of information across rows and columns. ZipHiC calculates the Potts normalising constant with an Approximate Bayesian Computation (ABC) sampler because otherwise it is computationally intractable. ZipHiC further classified contact counts into multiple components (“noise”, “signal” and “false signal”) and reduced false positives relative to other methods. However, it models the noise using a zero-inflated Poisson distribution, which might not accurately represent the noise in genomics data.

Here, we present a similar statistical approach to detect biologically significant chromatin interactions, using the zero-inflated Negative Binomial instead of zero-inflated Poisson distribution. The assumption of zero-inflated Negative Binomial distribution allows for improvement in the identification of each component by taking into consideration overdispersion that might occur within the components. We performed systematic analysis on real data, which shows improved performance over existing methods. Finally, we make this method available as an R/Bioconductor package called HiCPotts https://bioconductor.org/packages/HiCPotts/ that allows modelling of the noise using either the zero-inflated Negative Binomial or the zero-inflated Poisson distribution.

## MATERIALS AND METHODS

### HiCPotts model

We previously described a similar model using the zero-inflated Poisson distribution to model the noise and signal in (13). Here, we improved the model and assumed the zero-inflated Negative Binomial distribution.

#### Notations

Using the contact matrix between pairs of bins generated from Hi-C experiments, let *y*_*ij*_ denote the observed contact frequency between bin *i* and bin *j* in *N* total bins, and *D*_*ij*_ denote the genomic distance between bin *i* and bin *j*. Let *GC*_*ij*_ denote the average percentage of Guanine and Cytosine, *TE*_*ij*_ denote the average number of transposable elements (TEs), and *ACC*_*ij*_ denote the average DNA accessibility score in bins *i* and *j*. For convenience and readability, we denote the interaction pair of bins *i* and *j* as *s* = {*i, j*} and also *D*_*s*_, *GC*_*s*_, *ACC*_*s*_ and *TE*_*s*_ to denote the observation value for interaction *s*.

#### Mixture model for data

We assume that the contact frequency density of the *y*_*ij*_ comes from the following mixture model of *K* components, and the first component is a zero-inflated Negative Binomial distribution (*ZINB*) which represents the noise (see below), and the other components are modelled using the Negative Binomial (*NB*) distribution:

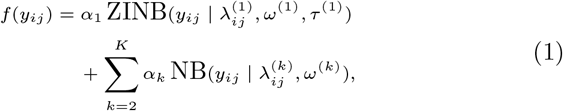

where

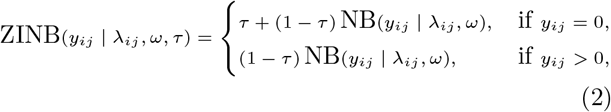

*τ* is the probability of extra zeros, *λ*_*ij*_ is the mean of the *k*th component, *ω* is the overdispersion parameter of the *k*th component. *α*_*k*_ is the unknown percentage of *k*th component subject to the constraint 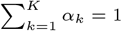. Since *ω* controls the degree of overdispersion, as *ω* → ∞ the equation approaches a Poisson (*λ*) model. A smaller value of *ω <* 1 implies that we have a case of overdispersion (15).

The above mixture model can be interpreted within a latent variable framework. We introduce the latent variable *z*_*ij*_ ∈ {1, 2, · · ·, *K*}, where *z*_*ij*_ = *k* indicates that *y*_*ij*_ follows the distribution of component *k*. Furthermore, 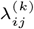 denotes the mean interaction between fragments *I* and *j* when it is from the *k*th component. Increasing the number of components *K* enhances the flexibility of the framework, allowing it to model diverse scenarios of interaction patterns.

Because the Hi-C (or micro-C) contact maps exhibit an excess of zero-counts and unequal mean-variance relationships, we assume that the noise follows a ZINB distribution rather than a ZIP distribution. Specifically, the ZINB distribution has mean (1 − *τ*)*λ* and variance 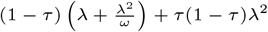. In addition, we corrected the sources of biases by modelling 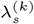 with *s* = {*i, j*}, *k* = 1, 2, · · ·, *K* as

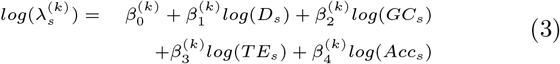

The unknown model parameters are denoted as **Θ** = (*τ, ω*, ***β***^(*k*)^, *k* = 1, · · ·, *K*) and the probability mass function for the observed data ***y***, given the latent variables ***z***, is denoted as *f* (***y***|**Θ, *z***).

To prevent label switching and preserve a consistent interpretation of mixture components across MCMC iterations, we impose an identifiability constraint by ordering components according to their baseline mean intensity. We define 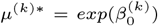 as the baseline mean contact frequency of the *k* − *th* component. To prevent label switching, we impose the ordering constraint *µ*^(1)∗^ *< µ*^(2)∗^ *<* … *< µ*^(*K*)∗^ in the prior. This yields stable component labels (“noise”, “signal”, “false signal”) across the chain and improves interpretability. We additionally assessed mixing and sensitivity in scenarios where component means are close, where ordering constraints can be less informative.

#### Potts Model

To capture the spatial dependence, our method employs a Hidden Markov Random Field (HMRF), which extends the widely known standard Hidden Markov Model (HMM). The HMRFs are widely applied across disciplines such as image analysis ((16)), gene expression data ((17)), and a population genetics study ((18)). Adapting the Potts model ((19)), a Markov random field-based method that provides a flexible way to model spatially dependent data, the prior distribution for the latent variable *z*_*s*_ is expressed as a function of its neighbours in a 2D grid. The latent variable ***z*** is written as

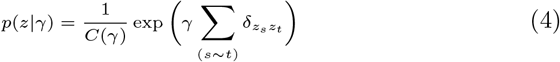

where 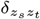 is the Kronecker symbol which takes the value 1 when *z*_*s*_ = *z*_*t*_ and 0 otherwise. *t* denotes the neighbouring fragment pairs of *s*, i.e. *s* ∼ *t* means *s* and *t* are neighbours in the Hi-C matrix. Our set of latent variables *z*_*ij*_ = *z*_*s*_ depends on the status of the neighbors of *s*, 𝒩 _*s*_ = {(*i* + 1, *j*), (*i* − 1, *j*), (*i, j* + 1), (*i, j* − 1)}. The neighbouring 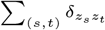 can be defined as the sum of the influence of neighbours of *s* controlled by a non-negative interaction parameter *γ*. When *γ* is 0, the latent variables behave independently. Higher values of *γ*, such as *γ* = 1, encourage spatial consistency, such that neighbouring bins are more likely to be placed in the same component. The normalizing constant term *C*(*γ*), also known as the partition function, is written as

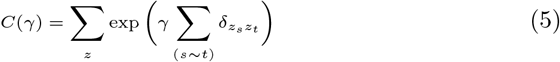

where ∑_*z*_ denotes the summation over *z*_*s*_ at all interactions *s* and it depends on the interaction parameter *γ*. The normalising constant term is computationally intractable in higher orders; therefore, methods such as the Approximate Bayesian Model (ABC) (20), a likelihood-free approach, can be used.

#### Approximate Bayesian Computation (ABC)

Given an observed dataset *Y* = (*y*_1_, *y*_2_, …, *y*_*n*_) that is associated with the models in equations (1) to (4), the ABC algorithm (20) used here is written as follows:

- Initialise parameters **Θ** and latent variables ***z*** and then repeat the following;
  1. Simulate an new value *γ*^∗^ from the prior distribution *π*_0_ (*γ*);
  2. Generate all latent variables *z*_*ij*_ from (4);
  3. Simulate a pseudo data set ***y***^∗^ from *f* (***y***|**Θ, *z***);
  4. Compute the distance (absolute or euclidean) *d* between the simulated data and the observed data;
  5. fix a tolerance *ϵ* or use an empirical quantile of *d*(*y*^∗^, *y*) which often corresponds to 1% quantile ((20))
  6. Accept *γ*^∗^ if the distance is less than *ϵ*, otherwise reject and start from step 1 again.
  7. Generate a parameter value **Θ** from the posterior distribution *π*_0_ (**Θ**)*f* (***y***|**Θ, *z***);

#### Bayesian Inference

A Bayesian framework is adopted, where the posterior distribution is proportional to the product of the prior and likelihood. Also, an Empirical Bayes approach is used to define a hierarchical prior structure, with hyper-parameters estimated from the data itself, thereby reducing sensitivity to prior mis-specification.

Further to the use of the Empirical Bayes approach, we also adopt the conventional Bayesian approach. For the conventional Bayesian approach, the prior for the parameter *β* follows the normal distribution. For example, 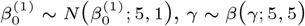 and *π*_0_ set to 0.5. See *Results* and *Supplementary Material*.

In this paper, we convert posterior means into component assignments using a fixed posterior-probability threshold: an interaction *s* is declared significant if *P* (*z*_*s*_ = 2(*signal*)|*y*) ≥ 0.90.

### Datasets and preprocessing

#### Drosophila dataset

Model performance was evaluated using the Hi-C map of Kc167 cell lines in *Drosophila* from (21). Raw data were downloaded and preprocessed with HiCExplorer following the set of parameters from (22; 23), with each pair of PE reads aligned separately to the *Drosophila melanogaster* (dm6) genome (24) using BWA-mem (25) (with options -t 20 -A1 -B4 -E50 -L0). HiCExplorer was then used to build and correct the contact matrices and detect enriched contacts (26) at 2 *Kb* bins resolution. The matrices were then exported in text format to be loaded in R.

We used DNaseI-seq data from (27) for DNA accessibility in *Drosophila* Kc167 cells data, and FlyBase (24) for TE annotation in *Drosophila*.

#### Human datasets

Hi-C and micro-C datasets from H1-hES cells from (28) were also analysed, following the same preprocessing pipeline used for the *Drosophila* dataset. We also aligned each pair to the human genome hg38 (29) using BWA-mem (25). Contact matrices were constructed using HiC-Explorer at 10 *Kb* resolution. HiCExplorer was also used to detect the enriched contacts (26).

In addition, DNaseI-seq were used to access DNA accessibility from the ENCODE consortium (30), with TE annotation sourced from RepeatMasker (31). For gene and TE annotations, we used UCSC genome browser annotations (32), while for enhancer annotations, we used Enhancer 2.0. (33).

### Comparison to other tools

In this paper, we compare our new method, HiCPotts using the Zero-Inflated Negative-Binomial to *(i)* ZipHiC (13), *(ii)* FastHiC (34), *(iii)* HiCExplorer (26) and *(iv)* Juicer (35). For enriched interactions detection, a JAVA implementation of FastHiC was applied using expected counts with values estimated by the HiCExplorer (26).

Then, HiCExplorer was run to generate the matrices, which are then corrected using the following values: (*i*) [−1.8, 5.0] for Hi-C in Kc167 cells, (*ii*) [−2.4, 5.0] for Hi-C in H1-hES cells and (*iii*) [−2.0, 5.0] for micro-C in H1-hES cells; see Supplementary (26). We generated the enriched contacts from the corrected matrix using `hicFindEnrichedContacts` tool with the observed over expected method (`--method obs/exp`) (26).

Furthermore, Juicer was used to generate enriched contacts by calling `dump` via its built-in tools. Specifically, we adopted the observed over expected method (oe) and Knight-Ruiz normalisation (KR) at 2 *Kb* resolution for the Hi-C data in Kc167 cells and at 10 *Kb* resolution for the Hi-C and micro-C data in H1-hES cells (35).

For the rest of the paper, we use HiCPotts or HiCPotts-ZINB to denote the use of the zero-inflated negative Binomial Distribution in the HiCPotts.

The R package used to perform the analysis can be downloaded from Bioconductor HiCPotts.

## RESULTS

We developed HiCPotts as an R/Bioconductor package to detect significant interactions from Hi-C, micro-C or Capture-C 3D chromatin data (see Figure 1). HiCPotts implements a Bayesian method based on the Hidden Markov Random Field HMRF and the Approximate Bayesian Computation ABC and uses a zero-inflated Negative Binomial distribution to model the noise, false signal and signal of the data (see *Materials and Methods*).

**Figure 1:**
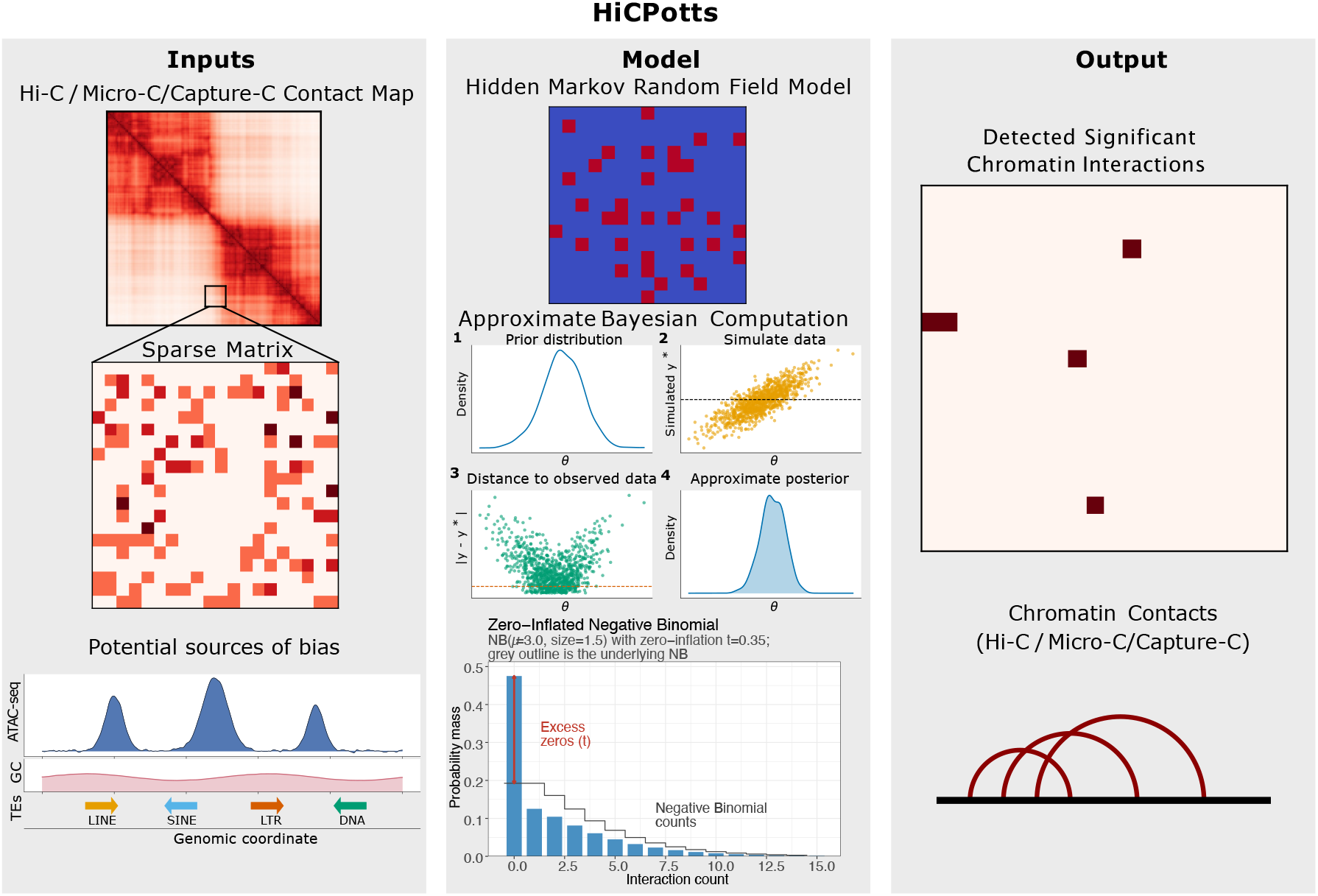
Workflow of HiCPotts.

### Identification of significant interactions in high resolution Hi-C data in *Drosophila melanogaster*

First, we evaluated the significant interactions detected by HiCPotts and compared them to ZipHiC a similar method that used zero-inflated Poisson distribution (thus, not modelling the over-dispersion) and FastHiC another, hidden Markov random field based method that does not model explicitly the sources of bias (such as DNA accessibility, GC content or Transposable Elements). Figure 2A compares the significant interactions detected by HiCPotts (ZINB), ZipHiC, and FastHiC within a region of *Drosophila* chromosome 2L. When HiCPotts uses an empirical data-driven prior, FastHiC identifies the largest number of interactions, followed by ZipHiC, while HiCPotts detects the smallest set (see Figure 2A). Despite this difference in total counts, all HiCPotts significant interactions are a subset of the overlap between ZipHiC and FastHiC. Our results show that adding sources of biases leads to losing approximately 3K interactions and further replacing the zero-inflated Poisson distribution with a zero-inflated Negative Binomial leads to losing an additional 17K significant interactions, with only approximately 5K interactions remaining detected by all three methods. Overall, we find that the ZINB is most stringent in detecting significant interactions and has a lower rate of false positives.

**Figure 2:**
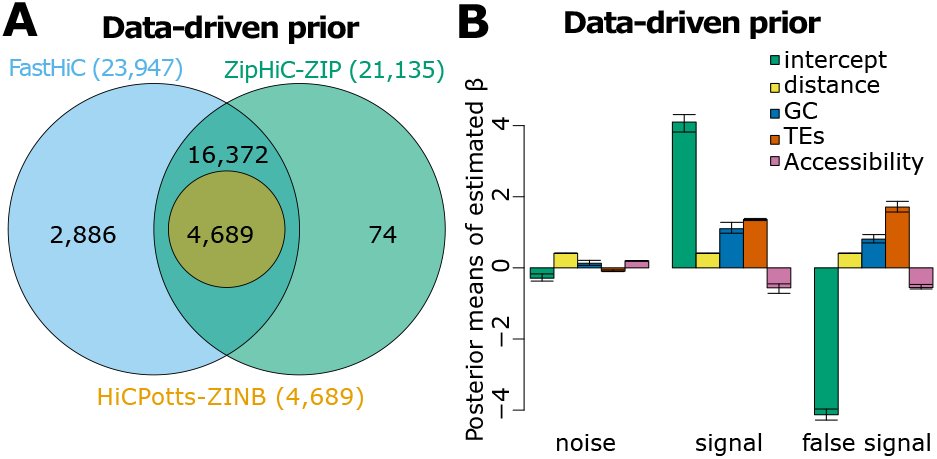
Comparison between HiCPotts (using ZINB and data-derived priors), ZipHiC and FastHiC. *(A)* Venn Diagram showing the comparison between HiCPotts (ZINB and data-derived priors), ZipHiC (ZIP) and FastHiC. The signficant interactions were detected on a subregion of chromosome 2L in *Drosophila* Kc167 cells. We considered that two significant interactions detected by the different tools are common if both anchors overlap fully, that is, the start and end of an anchor in one pair matches the start and end of the corresponding anchor in the other pair. The parameters for detecting the significant interactions can be found in the *Materials and Methods* section. *(B)* Posterior means of our estimated *β*s in the case of data derived priors. We computed the posterior means as shown in equation 3 for noise, signal and false signal components. The 95% credible intervals are shown.

A similar pattern is observed in Figure S1A, where HiCPotts is run with a user-defined prior. The number of HiCPotts-specific interactions changes only modestly between the two prior settings; see Figure S1B. 3, 560 (64.5%) were recovered under both, corresponding to 81.1% of the user-defined prior set and 75.9% of the data-driven set. This substantial overlap indicates that the core set of significant interactions identified by HiCPotts-ZINB is robust to prior specification and is driven primarily by the likelihood rather than the prior. The remaining 1, 961 prior-specific interactions (832 user-defined only, 1, 129 data-driven only) are concentrated near the posterior-probability decision boundary (*P* (*z*_*s*_ = *signal*|*y*) ≈ 0.90) and therefore reflect the expected sensitivity of any threshold-based classifier to small shifts in the posterior. The consistent and extensive overlap between HiCPotts and the other two methods across both sets of priors shows that HiCPotts with ZINB recovers a stringent subset of biologically meaningful 3D interactions.

To better understand the contributions of different components to the model and identify sources of bias, we analysed the posterior means of our estimated *β*s of the noise, signal and false signal for the data-derived priors; see Figure 2B. Negative values indicate that the corresponding component (noise, signal or false signal) is negatively correlated with the different features, while positive values indicate that the components are positively correlated with the different features. First, we observe that the *β* value for TEs on the false signal is high, indicating that regions with a higher number of TEs are more likely to be labelled as false signal by the HiCPotts model. Accessibility content is anticorrelated with the false signal and true signal. This means that some regions with lower accessibility tend to be labelled as false signal, while other regions with lower accessibility tend to be labelled as true signal by the model. Thus, the model is detecting a large number of contacts between silenced parts of the chromatin, either constitutive heterochromatin or Polycomb regions, and labels some of these as true signals while others as false signals. Polycomb regions have been shown to contribute to 3D contacts in Drosophila (36), supporting that some subset of the inaccessible parts of the genomes would display a high mean signal. Finally, GC content is lowly positively correlated with true signal and false signal, indicating potential bias from sequencing that affects the Hi-C data with regions with high GC content.

### Identification of significant interactions in medium resolution Hi-C data micro-C data in human ES cells

Next, we evaluated the performance of the method to identify significant interaction in Hi-C and micro-C data in human ES cells. In particular, we focused on the 60 − 70 *Mb* region of human chromosome 8 and used a 10 *Kb* resolution to compare HiCPotts (ZINB) to ZipHiC (using zero-inflated Poisson distribution) and two popular tools, HiCExplorer (26) and Juicer (35). Our results showed that all tools detect approximately 19*K* significant interactions for both Hi-C and micro-C; see Figure 3A-B. As observed previously, HiCPotts detects significantly less interactions that are a subset of the common interactions identified by the three other methods.

**Figure 3:**
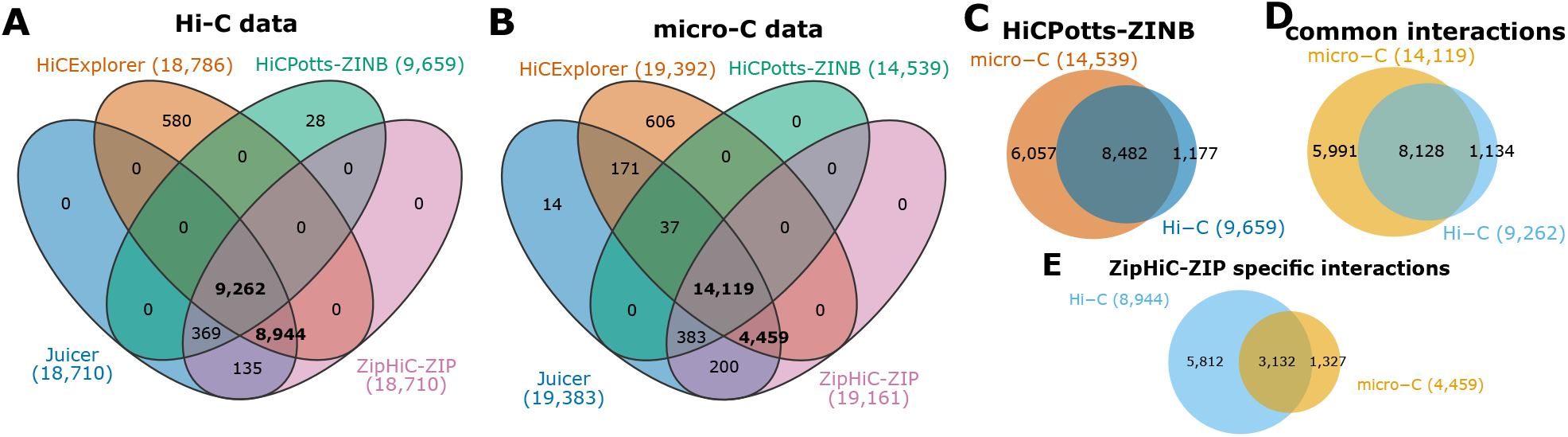
Significant interactions identified by HiCPotts and other methods on micro-C and Hi-C data in human ES cells. We consider a region of human chromosome 8 (60-70Mb). Two interactions detected by the different tools are common if both anchors overlap fully. The parameters for detecting the significant interactions can be found in the *Materials and Methods* section. *(A-B)* Interactions detected by the four methods in the *(A)* Hi-C data and *(B)* micro-C data. *(C)* Overlap of the interactions detected by HiCPotts in the Hi-C and micro-C data. *(D-E)* Overlap of the interactions detected in the Hi-C and micro-C data by *(D)* both HiCPotts and ZipHiC and *(E)* ZipHiC only.

Interestingly, we observe that HiCPotts detects approximately half of the interactions detected by other methods in the Hi-C data and approximately three-quarters in the micro-C data. To further dissect the differences between Hi-C and micro-C data, we compared the overlap of the significant interactions detected by HiCPotts in the Hi-C and micro-C data. Our results showed that almost all significant interactions detected in the Hi-C data are also detected in the micro-C data, but HiCPotts identified 6000 additional interactions in the micro-C; see Figure 3C. In micro-C, the chromatin is fragmented to mono-nucleosomes using micrococcal nuclease (MNase), which increases fragment density. These results indicate that although micro-C recovers many of the Hi-C interactions, it also identifies a substantially larger set of unique significant interactions, likely reflecting the increased resolution and denser fragmentation achieved through MNase-based mono-nucleosome fragmentation compared to restriction-enzyme fragmentation in Hi-C.

We labelled as ZipHiC specific interactions, interactions that are not detected in the Hi-C or micro-C data by HiCPotts but are detected by ZipHiC, Juicer and HiC-Explorer 3D-E. In the case of ZipHiC specific interactions, we find the opposite, namely that approximately 6000 interactions are only detected in the Hi-C data, and 3000 are detected in both Hi-C and micro-C data. Given that micro-C data has increased resolution and denser fragmentation suggests that some of these interactions that HiCPotts does not identify are likely false positives.

We further investigated the false signal interactions detected by HiCPotts-ZINB and found that 1, 671 were identified as being significant by Juicer, HiCExplorer and ZipHiC-ZIP in the micro-C data and 414 in the Hi-C data, indicating that a small subset of the contacts recovered by the other methods is being flagged as biased background by our model; see Figure S2. Consistent with the enrichment of ZipHiC-ZIP-specific calls for short-range contacts and their correlation with DNA accessibility and TE content, these shared false-signal interactions likely reflect uncorrected biases rather than genuine 3D contacts, supporting the advantage of overdispersion-aware modelling for distinguishing true signal from systematic noise.

Similarly, as in the case of the *D. melanogaster* Hi-C data, we analysed the posterior means of our estimated *β*s of the noise, signal and false signal for both micro-C and Hi-C data; see Figure 4. Interestingly, we observe that the DNA accessibility has a high *β* for the false signal component in the Hi-C data. This indicates that regions with higher accessibility are more likely to be labelled as false signal by HiCPotts model in the Hi-C data. The opposite is observed in the micro-C data, where more accessible regions are less likely to be labelled as false signal. In micro-C data, TE content seems to be correlated with false signal and noise and negatively correlated with the true signal. This means that regions with the highest TE content tend to be labelled more as false signal or noise and less as true signal by the model. Finally, GC content is lowly positively correlated with noise, true signal and false signal in both Hi-C and micro-C data. This suggests a potential modest bias from sequencing that affects both Hi-C and micro-C data similarly.

**Figure 4:**
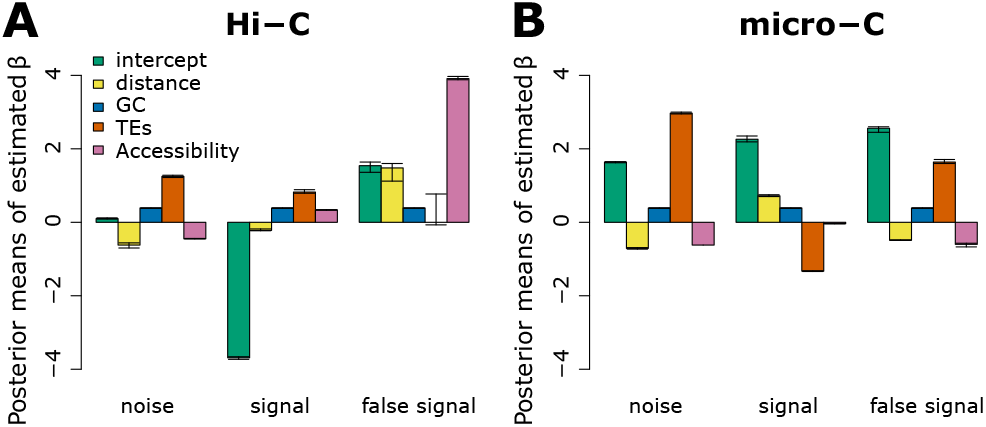
Posterior means of our estimated βs. We computed the posterior means as shown in equation 3 for noise, signal and false signal components of human Chromosome 8, region 60, 000, 000 : 70, 000, 000 for *(A)* Hi-C and *(B)* micro-C data. The 95% credible intervals are shown.

We further characterised the significant interactions detected by all methods (common interactions) and the ones not identified by HiCPotts but identified by all other methods (ZipHiC-ZIP specific). Figure 5A-D categorises the interactions according to functional annotations. For Hi-C data, HiCPotts-ZINB detects a higher proportion of interactions linking genes and intergenic regions (including TEs or regulatory regions) or promoters, while ZipHiC-ZIP specific interactions display a higher proportion of interactions between genes and enhancers (see Figure 5A-B). Similar results are observed for the micro-C data (see Figure5C-D). Interestingly, when investigating the distance between the two interacting regions, we found that HiCPotts-ZINB detects preferentially only long-range interactions (≥ 100 *Kb*), while ZipHiC-ZIP detects both long (≥ 100 *Kb*) and short (*<* 100 *Kb*) range interactions. Given that the resolution of the data is 10 *Kb*, interactions within 100 *Kb* distance are separated by less than 10 bins. Thus, these short-range interactions are more likely to be false positives.

**Figure 5:**
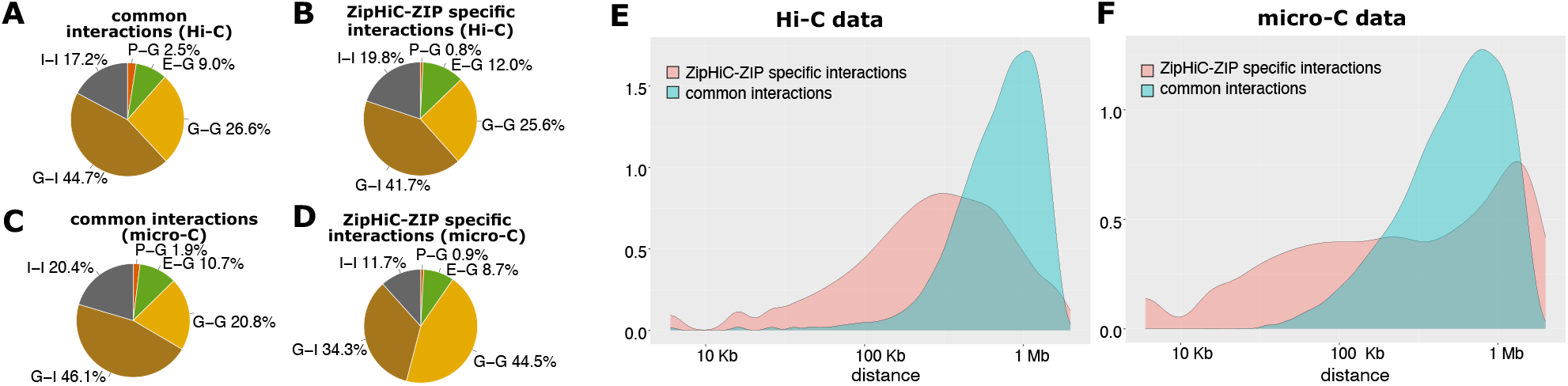
Characterisation of common and ZipHiC specific interactions in Hi-C and micro-C. *(A–D)* Distribution of genome annotation for the interactions identified in the 60∽70*Mb* region of human chromosome 8 by Hi-C and micro-C. Annotation of the regions linked by an interaction considers: (P) promoters (up to 1*kb* upstream of TSS), (E) enhancers, (G) genes, TE transposable elements, and (I) Intergenic. *(E–F)* Distance between the two interacting regions for the *(E)* Hi-C and *(F)* micro-C data. We considered separately common (detected by HiCPotts-ZINP) and ZipHiC specific interactions.

## DISCUSSION

In this paper, we introduced a new R/Bioconductor package, HiCPotts, that can detect significant interactions in Hi-C, micro-C or capture-C data. HiCPotts is a more comprehensive version of the ZipHiC method (13) and can model contact frequencies using a Zero-Inflated Negative Binomial (ZINB) distribution, a Zero-Inflated Poisson (ZipHiC) distribution, a Poisson distribution, and a Negative Binomial (NB) distribution. HiCPotts incorporates a Potts-model hidden Markov random field (HMRF) prior to account for spatial dependencies in the Hi-C contact matrix, similar to (13), but with improved variance modelling for overdispersion. Similar to ZipHiC (13), we adopt a likelihood-free ABC approach to overcome the intractable normalising constant of the Potts model and allow a three-component mixture (K = 3). This allows identification of false signals, along with noise and the true signal, in the data when present.

To investigate the most accurate distribution to model Hi-C or micro-C data, we model the contact frequencies using a Zero-Inflated Negative Binomial (ZINB) distribution, which captures the overdispersion characteristic of chromatin-contact data. Analyses of real datasets demonstrate that ZINB distribution achieves improved separation of noise and signal compared to existing tools. In particular, using the ZINB, we detect only a subset of the interactions detected by ZIP, namely approximately half for Hi-C and three-quarters for micro-C.

The noticeable difference in the number of significant interactions detected by the HiCPotts (ZINB) and the ZipHiC reflects the distinct statistical assumptions underlying the methods. HiCPotts (ZINB), which models the background using a zero-inflated Negative Binomial distribution and explicitly accounts for overdispersion, detects fewer significant interactions than ZipHiC, which uses a zero-inflated Poisson model. The HiCPotts (ZINB) model allows for a larger variance in the noise component, which leads to moderately high contact counts being accounted as part of a highly variable background rather than being attributed to the signal component. In contrast, the Poisson-based ZipHiC model underestimates the variance, making the same counts appear more extreme, and thus more likely to be classified as signal. This leads to a more conservative set of calls for HiCPotts (ZINB), enriched for robust interactions, and a larger set of calls for ZipHiC that includes the same interactions as HiCPotts (ZINB), but also additional, more borderline counts.

The interactions detected by HiCPotts-ZINB display a higher proportion of genes to regulatory regions interactions (intergenic, promoters or enhancers) than the interactions detected only by ZipHiC-ZIP. Most importantly, HiCPotts-ZINB preferentially detects only significant interactions beyond 100 *Kb*, while the interactions detected only by ZipHiC-ZIP are mostly between 10 *Kb* and 100 *Kb*. Previous work has shown that *cis*-regulatory regions control gene activity within 50 *Kb*, while, for longer distances, 3D contacts might be required (37), although that might not always be the case (38). This suggest that these shorter range interactions detected only by ZipHiC-ZIP might not only be false positives, but also might not have a functional role if they do exist.

While investigating the mixture components of the HiCPotts-ZINB model in human ES cells, we identified a high correlation between DNA accessibility and false signal in the Hi-C data with regions with higher accessibility being more likely to be labelled as false signal by our model. In addition, we also found that regions with higher TE content tend to be labelled more as false signal or noise and less as true signal in micro-C data. These results highlight the importance of including known potential sources of biases when modelling Hi-C or micro-C data.

While HiCPotts models additional features, including the overdispersion effect, spatial structure and multiple mixture components, that are not captured by other methods, this comes at a computational cost. HiCPotts, specifically when using the ZINB, is particularly well suited for Capture-C or targeted regions of Hi-C or micro-C datasets, where the number of bins is smaller, and the spatial structure can be exploited fully without heavy runtime.

In conclusion, HiCPotts offers a flexible and powerful framework for identifying significant chromatin interactions in Hi-C or micro-C data. By accounting for multiple assumptions for the contact frequencies distribution, options for user-specified priors, jointly modelling overdispersion, spatial structure, and multiple latent components, it distinguishes noise from true signal more accurately than existing methods and provides insight into the sources and magnitudes of experimental biases. Our results demonstrate that HiCPotts is a robust tool for dissecting and understanding the complex structure of 3D genome architecture.

## Author’s contributions

I.G.O., A.H., H.D., Y.B., and N.R.Z. conceived and designed the experiments. I.G.O. performed the experiments and analysed the data. A.H., H.D., Y.B., and N.R.Z. supervised the work. I.G.O., and N.R.Z. wrote the paper. The authors read and approved the final manuscript.

## Acknowledgements

The authors acknowledge the use of the High Performance Computing Facility at the University of Essex and would like to thank Stuart Newman for his support.

I.G.O was supported by University of Essex. N.R.Z. was supported by the Wellcome Trust (grant 202012/Z/16/Z). The analysis was facilitated by access to the Ceres high-performance computing cluster at the University of Essex.

## References

1. Bonev, B. & Cavalli, G. Organization and function of the 3D genome. Nature Reviews Genetics 17, 661–678 (2016).

2. Lieberman-Aiden, E. et al. Comprehensive mapping of long-range interactions reveals folding principles of the human genome. Science 326, 289–293 (2009).

3. Hsieh, T.-H. et al. Mapping nucleosome resolution chromosome folding in yeast by micro-c. Cell 162, 108–119 (2015). URL https://www.sciencedirect.com/science/article/pii/S0092867415006388.

4. Goel, V. Y., Huseyin, M. K. & Hansen, A. S. Region capture micro-c reveals coalescence of enhancers and promoters into nested microcompartments. Nature genetics 55, 1048–1056 (2023).

5. Yaffe, E. & Tanay, A. Probabilistic modeling of hi-c contact maps eliminates systematic biases to characterize global chromosomal architecture. Nature Genetics 43, 1059 (2011).

6. Imakaev, M. et al. Iterative correction of hi-c data reveals hallmarks of chromosome organization. Nature Methods 9, 999 (2012).

7. Knight, P. A. & Ruiz, D. A fast algorithm for matrix balancing. IMA Journal of Numerical Analysis 33, 1029–1047 (2013).

8. Carty, M. et al. An integrated model for detecting significant chromatin interactions from high-resolution hi-c data. Nature communications 8, 15454 (2017).

9. Sahin, M. et al. Hic-dc+ enables systematic 3d interaction calls and differential analysis for hi-c and hichip. Nature communications 12, 3366 (2021).

10. Fang, T. et al. Enhancing hi-c contact matrices for loop detection with capricorn: a multiview diffusion model. Bioinformatics 40, i471–i480 (2024).

11. Jiang, Z., Peng, Z., Luo, Y., Bie, L. & Wang, Y. Autohic: a deep-learning method for automatic and accurate chromosome-level genome assembly. bioRxiv 2023–08 (2023).

12. Yang, R., Karbalayghareh, A. & Leslie, C. S. Hic2self: self-supervised denoising for bulk and single-cell hi-c contact maps. bioRxiv 2024–11 (2024).

13. Osuntoki, I. G., Harrison, A., Dai, H., Bao, Y. & Zabet, N. R. Ziphic: a novel bayesian framework to identify enriched interactions and experimental biases in hi-c data. Bioinformatics (Oxford, England) btac 387 (2022).

14. Xu, Z. et al. A hidden markov random field-based bayesian method for the detection of long-range chromosomal interactions in hi-c data. Bioinformatics 32, 650–656 (2016).

15. Dai, H., Bao, Y. & Bao, M. Maximum likelihood estimate for the dispersion parameter of the negative binomial distribution. Statistics & Probability Letters 83, 21–27 (2013).

16. Zhang, Y., Brady, M. & Smith, S. Segmentation of brain mr images through a hidden markov random field model and the expectation-maximization algorithm. IEEE Transactions on Medical Imaging 20, 45–57 (2001).

17. Wei, Z., Li, H. et al. A hidden spatial-temporal markov random field model for network-based analysis of time course gene expression data. The Annals of applied statistics 2, 408–429 (2008).

18. François, O., Ancelet, S. & Guillot, G. Bayesian clustering using hidden markov random fields in spatial population genetics. Genetics 174, 805–816 (2006).

19. Wu, F.-Y. The potts model. Reviews of Modern Physics 54, 235 (1982).

20. Beaumont, M. A., Zhang, W. & Balding, D. J. Approximate bayesian computation in population genetics. Genetics 162, 2025–2035 (2002).

21. Eagen, K. P., Lieberman Aiden, E. & Kornberg, R. D. Polycomb-mediated chromatin loops revealed by a subkilobase-resolution chromatin interaction map. Proceedings of the National Academy of Sciences (2017).

22. Chathoth, K. T. & Zabet, N. R. Chromatin architecture reorganisation during neuronal cell differentiation in Drosophila genome. Genome research 29, 613–625 (2019). URL https://www.biorxiv.org/content/early/2018/08/30/395822 http://www.ncbi.nlm.nih.gov/pubmed/30709849.

23. Chathoth, K. T. et al. The role of insulators and transcription in 3D chromatin organization of flies. Genome Research 32, 682–698 (2022). URL https://genome.cshlp.org/content/early/2022/03/30/gr.275809.121 https://genome.cshlp.org/content/early/2022/03/30/gr.275809.121.abstract.

24. dos Santos, G. et al. FlyBase: introduction of the Drosophila melanogaster Release 6 reference genome assembly and large-scale migration of genome annotations. Nucleic Acids Research 43, D690–D697 (2015).

25. Li, H. & Durbin, R. Fast and accurate long-read alignment with Burrows-Wheeler transform. Bioinformatics 26, 589–595 (2010).

26. Ramirez, F. et al. High-resolution TADs reveal DNA sequences underlying genome organization in flies. Nature Communications 9, 189 (2018). s41467-017-02525-w.

27. Kharchenko, P. V. et al. Comprehensive analysis of the chromatin landscape in Drosophila melanogaster. Nature (2010).

28. Krietenstein, N. et al. Ultrastructural details of mammalian chromosome architecture. Molecular Cell 78, 554–565.e7 (2020). URL https://www.sciencedirect.com/science/article/pii/S1097276520301519.

29. Schneider, V. A. et al. Evaluation of GRCh38 and de novo haploid genome assemblies demonstrates the enduring quality of the reference assembly. Genome Research 27, 849–864 (2017). URL http://www.genome.org/cgi/doi/10.1101/gr.213611.116.

30. Thurman, R. E. et al. The accessible chromatin landscape of the human genome. Nature 489, 75–82 (2012). URL http://www.nature.com/articles/nature11232.

31. Smit, H. R.. G. P., AFA. RepeatMasker Open-4.0. http://www.repeatmasker.org (2013-2015).

32. Casper, J. et al. The ucsc genome browser database: 2026 update. Nucleic Acids Research 54, D1331–D1335 (2026).

33. Gao, T. & Qian, J. Enhanceratlas 2.0: An updated resource with enhancer annotation in 586 tissue/cell types across nine species. Nucleic Acids Research 48, D58–D64 (2020). URL https://academic.oup.com/nar/article/48/D1/D58/5628925.

34. Xu, Z., Zhang, G., Wu, C., Li, Y. & Hu, M. Fasthic: a fast and accurate algorithm to detect long-range chromosomal interactions from hi-c data. Bioinformatics 32, 2692–2695 (2016).

35. Durand, N. C. et al. Juicer Provides a One-Click System for Analyzing Loop-Resolution Hi-C Experiments. Cell Systems 3, 95–98 (2017).

36. Ogiyama, Y., Schuettengruber, B., Papadopoulos, G. L.Chang, J.-M. & Cavalli, G. Polycomb-dependent chromatin looping contributes to gene silencing during drosophila development. Molecular Cell 71, 73–88.e5 (2018). URL https://www.sciencedirect.com/science/article/pii/S1097276518304076.

37. Guckelberger, P. et al. Cohesin-mediated 3d contacts tune enhancer-promoter regulation. bioRxiv (2024). URL https://www.biorxiv.org/content/early/2024/07/22/2024.07.12.603288. https://www.biorxiv.org/content/early/2024/07/22/2024.07.12.603288.full.pdf.

38. Benabdallah, N. S. et al. Decreased enhancer-promoter proximity accompanying enhancer activation. Molecular Cell 76, 473–484.e7 (2019). URL https://doi.org/10.1016/j.molcel.2019.07.038. Doi: 10.1016/j.molcel.2019.07.038.

