## Supplementary Material for "HiCPotts: An R/Bioconductor package to identify significant interactions in chromosome conformation capture data and model sources of biases"

### 1 Model for Data

The complete likelihood function of the unknown parameters $(\boldsymbol{\beta}, \boldsymbol{\alpha}, \boldsymbol{\tau})$ given the data $y$ and labels $z$ can then be written as

$$l(\vec{\boldsymbol{\beta}}, \vec{\boldsymbol{\omega}}, \boldsymbol{\alpha}, \boldsymbol{\tau} \mid \boldsymbol{y}, \boldsymbol{z}) \propto \prod_{i=1}^{n}\prod_{j=1}^{n}\Big\{ \Big[\alpha_1\Big(\tau\,\mathbb{I}\{y_{ij}=0\} + (1-\tau)\,\mathrm{NB}\Big(y_{ij}\,;\,\lambda_{ij}^{(1)},\,\omega^{(1)}\Big)\Big)\Big]^{\mathbb{I}(z_{ij}=1)} \prod_{k=2}^{K}\Big[\alpha_k\,\mathrm{NB}\Big(y_{ij}\,;\,\lambda_{ij}^{(k)},\,\omega^{(k)}\Big)\Big]^{\mathbb{I}(z_{ij}=k)}\Big\}, \tag{S1}$$

where $\mathrm{NB}\Big(y;\lambda,\omega\Big) = \frac{\Gamma(y+\omega)}{\Gamma(\omega)\,y!}\left(\frac{\omega}{\omega+\lambda}\right)^{\omega}\left(\frac{\lambda}{\omega+\lambda}\right)^{y}, \quad y \in \{0,1,2,\ldots\},$

The full posterior of $z$, $\vec{\beta}$ and $\gamma$ given $y_{ij}$ is

$$Pr(\boldsymbol{z}, \vec{\boldsymbol{\beta}}, \gamma|\boldsymbol{y}) \propto l(\vec{\boldsymbol{\beta}}, \boldsymbol{\alpha}, \boldsymbol{\tau}|\boldsymbol{y}, \boldsymbol{z})\,l(\boldsymbol{z}|\gamma)\pi_0(\gamma)\tau_0(\vec{\boldsymbol{\beta}}) \tag{S2}$$

where $l(\boldsymbol{z}|\gamma) = \frac{e^{\gamma\sum_{s\sim t}\delta_{z_s z_t}}}{\sum_{z_s} e^{\gamma\sum_{s\sim t}\delta_{z_s z_t}}}$ is the Potts model, $s$ is bin pair $i$ and $j$, and $t$ is the neighbours set of $s$; $(i-1, i+1, j-1, j+1)$.

In order to analyse our data and estimate our parameters, we make use of the Metropolis-within-Gibbs sampler and the Approximate Bayesian Computation (ABC), so the conditional posterior densities are needed.

#### 1.1 Conditional Posterior Density

The conditional posterior of $\tau$ is given as

$$Pr(\tau|\vec{\boldsymbol{\beta}}^{\mathbf{1}}, \boldsymbol{y}, \boldsymbol{z}) \propto \prod_{i=1}^{n}\prod_{j=1}^{n}[f^{(1)}(y_{ij};\boldsymbol{\beta}^{(1)})]^{I[z_{ij}=1]}\cdot\pi_0(\tau) \tag{S3}$$

The conditional posterior of $\beta$ is given by

$$Pr(\vec{\boldsymbol{\beta}}^{(\boldsymbol{k})}|\boldsymbol{y}, \boldsymbol{z}) \propto \prod_{i=1}^{n}\prod_{j=1}^{n}\prod_{k}^{K}[f^{(k)}(y_{ij};\boldsymbol{\beta}^{(k)})]^{I[z_{ij}=k]}\pi_0(\boldsymbol{\beta}^{(k)}) \tag{S4}$$

Based on the definitions of $f^{(k)}(y_{ij})$ and equation S4, the conditional posterior of $\beta^{(1)}$ for the noise (ZI) component can be rewritten as

$$\Pr(\vec{\boldsymbol{\beta}}^{(\mathbf{1})} \mid \boldsymbol{y}, \boldsymbol{z}) \propto \prod_{i=1}^{n}\prod_{j=1}^{n}\Big[\Big(\tau\,\mathbb{I}\{y_{ij}=0\} + (1-\tau)\,\mathrm{NB}\Big(y_{ij};\,\lambda_{ij}^{(1)},\,\omega^{(1)}\Big)\Big)\Big]^{\mathbb{I}(z_{ij}=1)} \exp\Bigg\{-\frac{\big(\beta^{(1)}-m^{(1)}\big)^2}{2\,(\sigma^2)^{(1)}}\Bigg\}. \tag{S5}$$

For the signal component, the conditional posterior of $\beta^{(k)}$ (for $k \geq 2$) based on the definition of $f^{(k)}(y_{ij})$ and equation S4 can be rewritten as

$$\Pr(\vec{\boldsymbol{\beta}}^{(\boldsymbol{k})} \mid \boldsymbol{y}, \boldsymbol{z}) \propto \prod_{i=1}^{n}\prod_{j=1}^{n}\Big[\mathrm{NB}\Big(y_{ij};\,\lambda_{ij}^{(k)},\,\omega^{(k)}\Big)\Big]^{\mathbb{I}(z_{ij}=k)} \exp\Bigg\{-\frac{\big(\beta^{(k)}-m^{(k)}\big)^2}{2\,(\sigma^2)^{(k)}}\Bigg\}. \tag{S6}$$

where $m^{(k)}$ is the mean and $\left(\sigma^{(k)}\right)^2$ is the variance for component $k$. Here $\mathrm{NB}(y;\lambda,r)$ denotes the negative binomial pmf parameterised by mean $\lambda$ and size $r$, and $\lambda_{ij}^{(k)} = \exp\left(x_{ij}^{\top}\beta^{(k)}\right)$ under the log link.

The conditional posterior of the NB size parameter ($\omega$) is given as

$$\Pr\Big(\omega^{(1)} \mid \vec{\boldsymbol{\beta}}^{(1)}, \tau, \boldsymbol{y}, \boldsymbol{z}\Big) \;\propto\; \prod_{i=1}^{n}\prod_{j=1}^{n}\Big[\,f^{(1)}\Big(y_{ij};\,\boldsymbol{\beta}^{(1)}, \omega^{(1)}, \tau\Big)\Big]^{\mathbb{I}[z_{ij}=1]} \cdot \pi_0(\omega^{(1)}). \tag{S7}$$

$$\Pr\Big(\vec{\omega}^{(k)} \mid \vec{\boldsymbol{\beta}}^{(k)}, \boldsymbol{y}, \boldsymbol{z}\Big) \;\propto\; \prod_{i=1}^{n}\prod_{j=1}^{n}\Big[\,f^{(k)}\Big(y_{ij};\,\boldsymbol{\beta}^{(k)}, \omega^{(k)}\Big)\Big]^{\mathbb{I}[z_{ij}=k]} \cdot \pi_0(\omega^{(k)}). \tag{S8}$$

To update the latent variable, the probability of an observation belonging to each component is calculated

$$Pr(z_s|\gamma, z_t, \boldsymbol{y}, \vec{\boldsymbol{\beta}}^{(\boldsymbol{k})}) \propto e^{\gamma\sum_{s\sim t}\delta_{z_s z_t}} f(y_{i,j}; \boldsymbol{\beta}^{(k)}) \tag{S9}$$

where $s$ is bin pair $i$ and $j$, and $t$ is the neighbours set of $s$; ($i-1$, $i+1$, $j-1$, $j+1$) and $f^{(k)}(y_{ij}; \vec{\boldsymbol{\beta}}^{(\boldsymbol{k})})$ is the likelihood of component $k$.

When the normalizing constant is introduced, equation S9 can be rewritten as

$$Pr(z|\gamma, z_t, \boldsymbol{y}, \vec{\boldsymbol{\beta}}^{(k)}) = \frac{e^{\gamma\sum_{s\sim t}\delta_{z_s z_t}}}{\sum_{z_s} e^{\gamma\sum_{s\sim t}\delta_{z_s z_t}}} \frac{f(y_{i,j}; \boldsymbol{\beta}^{(k)})}{f(y_{i,j}; \boldsymbol{\beta}^{(k)})} \tag{S10}$$

where $s$ is bin pair $i$ and $j$, and $t$ is the neighbour(s).

The conditional probability of $\gamma$ in the Potts model is given as

$$Pr(\gamma|\boldsymbol{y}, \boldsymbol{z}, \vec{\boldsymbol{\beta}}) = \frac{\exp\{\gamma\sum_{s\sim t}\delta(z_s z_t)\}\pi_0(\gamma)}{\sum_{z_s}\exp\{\gamma\sum_{s\sim t}\delta(z_s z_t)\}\pi_0(\gamma)} \tag{S11}$$

Algorithm 1: ABC

**repeat**
    Select the initial value of $\gamma^0$;
    $m = 0$
    **for** $i = 1 : N$ **do**
        Compute a new $y^*$ based on the Potts model and updated $\vec{\beta}$ from Algorithm (2)
        Compute the distance $d(S(y^*), S(y))$
        Select $\epsilon$ using $1\%$ empirical quantile of the distance or other chosen distance metric
        **if** $d(S(y^*), S(y)) < \epsilon$ **then**
            $\gamma_{m+1} = (\gamma_m)^*$
        **end ifend if**
    **end forend for**
**until** enough MCMC steps have been simulated

[1]

Algorithm 2: Metropolis-within-Gibbs sampler

**procedure**
    Initialization, select initial value,$z^0, \gamma^0, \tau^0, \omega^0, \vec{\beta}^0$;
    **repeat**
        **for** $i = 1$ to $n$, $j = 1$ to $n$ **do**
            Update $z_{ij}$ using (S11)
        **end forend for**
        Update $\vec{\beta}$ from posterior in (S5)and (S6)
        Update $\tau$ using (S3)
        Update $\omega$ from posterior in (S7) and (S8)
        Update $\gamma$ using Algorithm (1)
    **until** enough MCMC steps have been simulated;
**end procedure**

The equations for $\vec{\beta}$'s, $\vec{\omega}$'s and $z$ as given above in equations S3, S5, S6, S7 and S8 is computationally easier to simulate using the Metropolis-Hastings-within-Gibbs sampler. Equation S11 is computationally intractable as the interaction parameter $\gamma$ involves the evaluation of the partition function and cannot be simulated directly using the Gibbs sampler or the Metropolis-Hastings sampler.

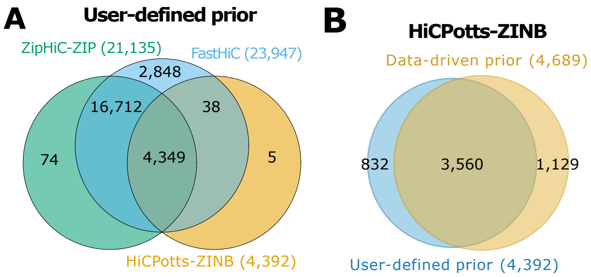


Figure 1: *Comparison between HiCPotts (using ZINB and user-derived priors), ZipHiC and FastHiC. (A)* Venn Diagram showing the comparison between HiCPotts (ZINB with user-derived priors), ZipHiC (ZIP) and FastHiC. The signficant interactions were detected on a subregion of chromosome $2$L in *Drosophila* Kc167 cells. We considered that two significant interactions detected by the different tools are common if both anchors overlap fully, that is, the start and end of an anchor in one pair matches the start and end of the corresponding anchor in the other pair. The parameters for detecting the significant interactions can be found in the *Materials and Methods* section. *(B)* The overlap between the significant interactions detected by HiCPotts (using ZINB) using the two sets of priors (data-driven or user-defined).

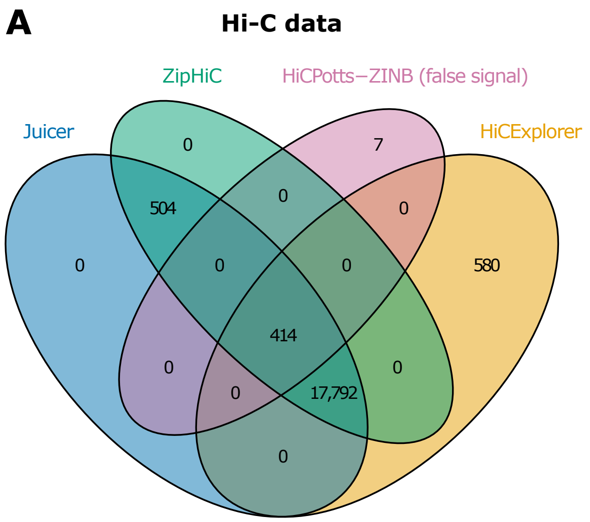


Figure 2: *Significant interactions identified by HiCPotts as false signal and as true signal by other methods on micro-C and Hi-C data in human ES cells.* We consider a region of human chromosome 8 (60-70Mb). Two interactions detected by the different tools are common if both anchors overlap fully. The parameters for detecting the significant interactions can be found in the *Materials and Methods* section. *(A-B)* Interactions detected by the four methods in the *(A)* Hi-C data and *(B)* micro-C data.

Algorithm 1 shows the Approximate Bayesian Computation (ABC) approximate steps and Algorithm 2 shows the Metropolis-within-Gibbs steps used in this paper to update our parameters.
